# A keeper of many crypts: the parasitoid *Euderus set* manipulates the behavior of a taxonomically diverse array of oak gall wasp species

**DOI:** 10.1101/516104

**Authors:** Anna K. G. Ward, Omar S. Khodor, Scott P. Egan, Kelly L. Weinersmith, Andrew A. Forbes

## Abstract

Parasites of animals and plants can encounter trade-offs between their specificity to any single host and their fitness on alternative hosts. For parasites that manipulate their host’s behavior, the complexity of that manipulation may further limit the parasite’s host range. The recently described crypt-keeper wasp, *Euderus set*, changes the behavior of the gall wasp *Bassettia pallida* such that *B. pallida* chews an incomplete exit hole in the side of its larval chamber and “plugs” that hole with its head. *E. set* benefits from this head plug, as it facilitates the escape of the parasitoid from the crypt after it completes development. Here, we ask whether this behavioral manipulator is limited to *Bassettia* hosts. We find that *E. set* attacks and manipulates the behavior of at least six additional gall wasp species, and that these hosts are taxonomically diverse. Interestingly, each of *E. set*’s hosts has converged upon similarities in their extended phenotypes: the galls they induce on oaks share characters that may make them vulnerable to attack by *E. set*. Behavioral manipulation in this parasitoid system may be less important to its host range than other dimensions of the host-parasitoid interaction, like the host’s physical defenses.

## 1. Introduction

Evolutionary biologists have proposed that trade-offs present limits to adaptation [1], as resources are finite and optimization of all traits at once is impossible. This is most clear when energy needs to be allocated to multiple competing functions. For example, when energy is dedicated towards growth, it may not be available for reproduction [2]. One common trade-off in nature is the observation that when parasites adapt to attack one group of hosts, those traits are often maladaptive for other hosts. This trade-off has been used to explain, in part, the tendency for parasites to be more specialized (as opposed to generalized) and only feed on a subset of available hosts (e.g., herbivorous insects on plant hosts or blood feeding insects on animal hosts) [3].

The ability of parasites to control their host’s behavior may be a trait especially likely to result in increased specialization and reduced fitness outcomes on alternative hosts. Parasite manipulation of host behavior is a phenomenon in which a parasite changes the activities of its host in ways that are good for the parasite, but are typically bad for the host [4–7]. Manipulation of host behavior may involve parasites producing neuroactive compounds, influencing connections between the immune and nervous systems, or inducing genomic/proteomic changes in the host, all with the goal of producing changes in host behavior that favor the parasite [8]. Such intimate control of host physiology and behavior might reasonably be expected to limit the number of host species a parasite can manipulate.

What do we know about the relationship between how parasites manipulate host behavior and parasite host range? An analysis by Fredensborg [9] examined this question in the context of trophically transmitted parasites (i.e., parasites that need their current host to be consumed by the next host in the parasite’s life cycle [10]) that manipulate host behavior. This analysis revealed no difference in host specificity between parasites whose manipulation was categorized as “simple” (specifically, they altered host activity levels to make it more susceptible to predation by the next host on the parasite’s life cycle), and those whose manipulation was categorized as “complex” (in which parasites increased the amount of time spent by hosts in microhabitats frequented by the next host). Fredensborg’s [9] work is a valuable first look at the relationship between manipulation of hosts and host range, though we note that binning manipulated behaviors in this way may obscure the multidimensionality of manipulation [11] and the same behavior can be changed through multiple mechanisms which may differ in how complex they are for the parasite to achieve.

The trophically transmitted parasites analyzed by Fredensborg [9] differ in critical ways from other types of manipulators. For example, the most common outcome of manipulation by trophically transmitted parasites is an increased likelihood that the host and its parasite are consumed by the right host at the right time [12,13], while parasitoids (i.e., parasites that need to kill their hosts in order to complete their life cycle [14]) may manipulate hosts to reduce the likelihood that their offspring are attacked by various natural enemies [15]. These two categories of parasites also tend to differ taxonomically (trophically transmitted parasites are often helminths, while parasitoids are often insects), and in size of the parasite relative to its host [14]. These major differences between types of manipulators suggests that analyses are needed for each manipulator type to explore the generality of rules associated with limits on the numbers of hosts a parasite can manipulate. One reason these types of analyses are rare is because they require careful work, including detailed natural history, specialized taxonomy, surveys of museum records, and challenging field work, searching for evidence of manipulation in multiple different host species, and careful work confirming the identity of the parasite infecting each of these hosts. Data such as these are critical if we are to understand the evolution of manipulation, and limits on the number of hosts a parasite can manipulate. Here, we help fill this gap by studying host specificity in a parasitoid that manipulates the behavior of cynipid gall wasps.

Two recent papers [16,17] describe the discovery and life history of the parasitoid “crypt keeper wasp” (*Euderus set*), so named because of its parasitism and behavioral manipulation of the “crypt-gall wasp” (*Bassettia pallida*) in live oaks (*Quercus virginiana & Q. geminata*). Oviposition by a *B. pallida* female into a young stem induces the formation of a swollen internal gall (known as a “crypt”) inside the stem, where the developing larval wasp will then feed and grow. Unparasitized adult *B. pallida* later chew a small exit hole in their gall and fly away. However, when parasitized by *E. set, B. pallida* chew significantly smaller exit holes, do not (or cannot) leave the gall, and die with their heads blocking (or “plugging”) the exit hole. *Euderus set* then feeds on the now disabled body of the host wasp, and, upon maturing into an adult wasp, chews through the “head plug” and exits the gall (Figure 1). Because *B. pallida* manipulates oaks to develop galls in their young branches, *E. set* is a “hypermanipulator” – a rarely quantified phenomena in which a parasite manipulates a parasite that itself manipulates its host. Though many species of *Euderus* have previously been described as parasitoids of a taxonomically diverse series of insect hosts [18], *E. set* represents the first definitive example of behavioral manipulation in this genus.

**Figure 1.**
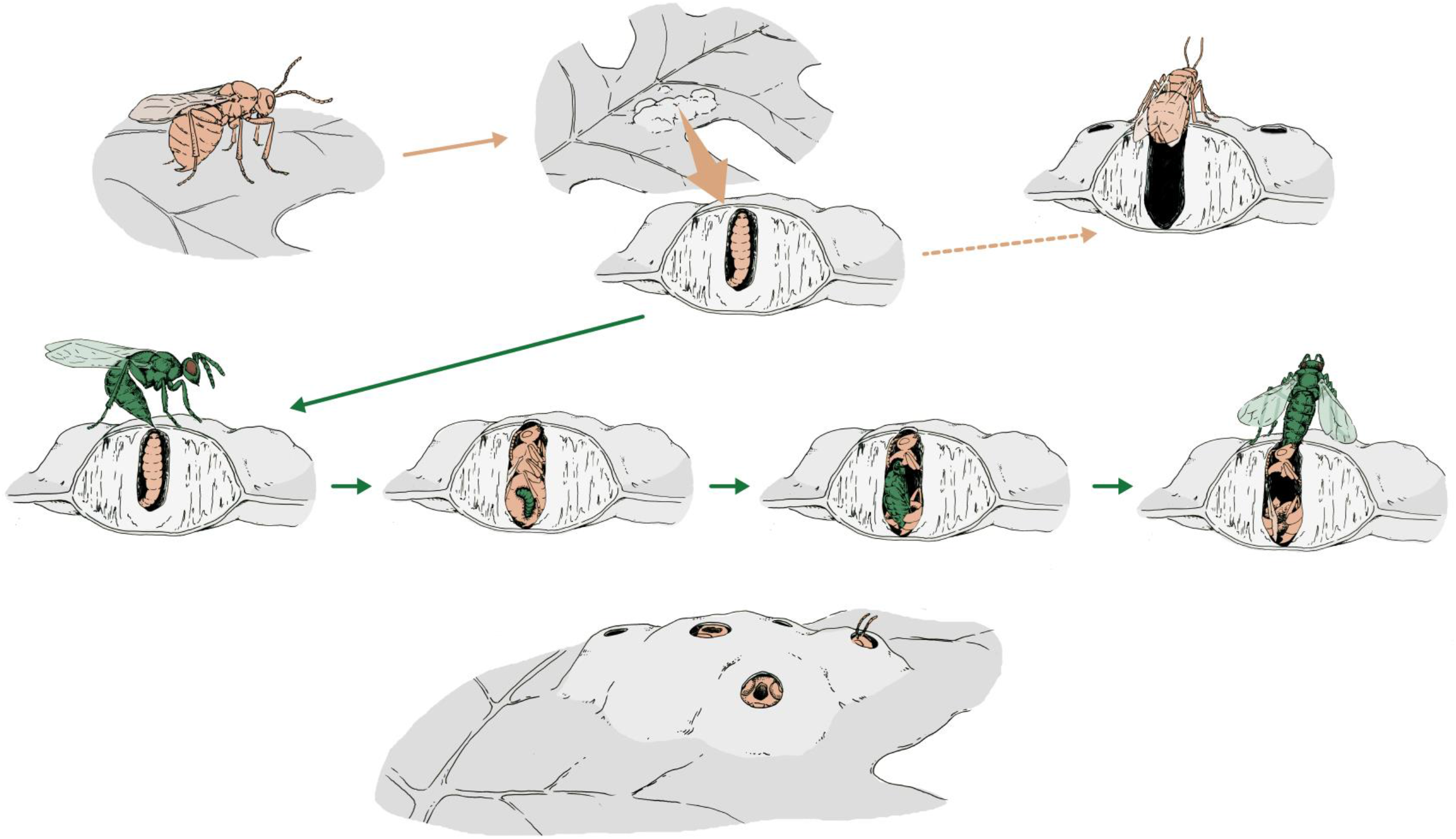
Life cycle of an oak gall wasp (in this case, *Callirhytis quercusmodesta*; pink), and the crypt keeper wasp, *Euderus set* (green). Artwork by Mona Luo.

The parasitoid communities of most oak gall wasps are understudied or unknown such that it is premature to assert that the manipulation of *B. pallida* by *E. set* is a unique relationship. Indeed, Weinersmith et al. [17] report additional examples of “head plugging” in the gall wasp *Bassettia ligni* on the host plants *Q. lobata* and *Q. douglasii*. These examples were from museum specimens, which were not destructively sampled to confirm the presence of *Euderus* within the gall. They also report evidence of head-plugging in an unidentified cynipid on *Q. nigra* in southeast Texas [17], that has since been keyed to genus *Bassettia* (possibly *B. aquaticae*; A.A.F. unpublished data). Further, our own parasitoid rearing studies have yielded several unidentified *Euderus* from *non-Bassettia* galls (A.K.G.W. and A.A.F., unpublished data). These observations raise at least two questions about *Euderus*. First, are these other unidentified *Euderus* also members of the species *E. set*, or do different *Euderus* species attack different species of North American oak gall wasps? And second, is behavioral manipulation of oak gall wasps limited to the *Euderus* that attack *Bassettia* hosts, or do other *Euderus* associated with oak gall wasps also manipulate their hosts’ behavior?

The above two questions are not mutually exclusive, such that four outcomes are possible for this study. First, we may find a “many manipulating specialists” scenario, in which many *Euderus* species attack North American oak gallers and each induces head plugs in its respective host. Second, we might find many “many specialists, few manipulators,” i.e., that several *Euderus* species may each attack one or a few gall wasp species, but only *E. set* – or a subset of species – induce the head-plugging phenotype. Third, *E. set* may be a lone “master manipulator,” attacking and inducing head plugs in several hosts. Or fourth, *E. set* may a “contingent manipulator,” attacking many oak gall wasp species but only inducing the head plugging phenotype in *Bassettia* galls. We consider “head-plugging” to be a relatively simple manipulation, as it requires the host to initiate a behavior it would have performed in its uninfected state, yet the parasitoid stops the behavior before completion. However, without knowing the mechanism through which this manipulation is achieved, it is difficult to favor one of the proposed hypotheses above the others.

Here, we evaluate support for these four scenarios by studying the identity and behaviors of *Euderus* reared from six oak gall wasp host species in seven species of oak tree across a wide geographic range. In doing so, we address a critical question and a gap in our current knowledge regarding how manipulation of insect behavior by parasitoids is (or is not) translatable across disparate hosts, and provide critical information for future analyses examining constraints on the number and type of hosts a parasitoid can manipulate.

## 2. Methods

### (a) *Collections and discovery of new* Euderus / *galler associations*

From August 2015 to August 2018, we collected more than 23,000 galls from a variety of oak species and locations, with a focus on maximizing the diversity of gall wasp species and on collecting mature galls that were most likely to have been parasitized. All galls were North American, and most collections (~60%) were made in Midwestern states, but at least some collections extended farther afield, including to e.g., New England, North Carolina, and Texas (Supplementary Table 1). We placed oak galls of the same gall wasp species, tree host, collection date, and location in individual cups in an environmental chamber (SANYO Electric Co. Ltd, Osaka, Japan) that approximated the average day and night temperatures and light and dark cycles in Iowa City, IA, with the exception that we used a minimum temperature of 5°C during winter months. We checked the incubator daily for the presences of emergent animals and placed new emergences in 95% ethanol at −80°C. Emergent animals included gall wasps and a large diversity of parasitoids, inquilines, and hyperparasitoids, which we identified to family or genus using taxonomic keys found in [19] and [20]. We identified *Euderus* to species using [18] and Egan et al. [16]. Most gall wasp species are identifiable by their gall’s morphology, such that keying out emerging adult gall wasps to species was largely unnecessary.

### (b) MtCOI sequencing and assessment of species status

When *Euderus* emerged from galls, we extracted DNA from representative samples using a CTAB/PCI method based on Chen et al. [21]. We PCR amplified the mtCOI region using the following primers: COI_PF2 5’ ACC WGT AAT RAT AGG DGG DTT TGG DAA 3’ and COI_2437d 5’ CGT ART CAT CTA AAW AYT TTA ATW CCW G 3’ [22]. For four samples sequenced in 2016, we used the following primers: LEP F 5’ TAA ACT TCT GGA TGT CCA AAA AAT CA 3’ and LEP R 5’ ATT CAA CCA ATC ATA AAG ATA TTG G 3’ [23]. We then used EXO1 and SAP to clean PCR products before Sanger sequencing in both forward and reverse directions on an ABI 3720 DNA Analyzer (Applied Biosystems, Foster City, CA) in the University of Iowa’s Roy J. Carver Center for Genomics. We aligned reverse and forward sequences in the program Geneious 8.18 (Biomatters Ltd, Newark, NJ). Most of the resulting fragments were 701bp in length, while the four sequenced in 2016 were 652bp, with 464 bp of overlap between the two fragment types. We generated a multiple alignment of these new *Euderus* mtCOI sequences and two known *E. set* sequences [16] (Supplementary Table 2). For outgroups, we used sequences from NCBI from four other species in the same subfamily, including *Euderus albitarsus* and *Euderus cushmanii*, introduced and native North American (respectively) parasites of Lepidopteran pupae [18]. We inferred a Bayesian tree using Mr.Bayes 3.2.6 [24] and also directly compared the percentage similarity among sequences.

### (c) Observations of Euderus behavior and emergence phenology

We chose one galler species for a focused study of the natural history, behavior, and phenology of *Euderus* host manipulation in a *non-Bassettia* gall. From June 15^th^ to July 20^th^, 2018, we collected *Callirhytis quercusmodesta* galls weekly from a single, heavily infested pin oak (*Quercus palustris*) tree in Iowa City, IA. Galls of *C. quercusmodesta* manifest as parenchymal thickenings that project on both sides of the leaf. Each gall contains multiple larval chambers ranging from a dozen to upwards of a hundred (Figure 2A). Upon collection, we cut each multi-chambered gall from its leaf, assigned it a number, photographed it from above using a Canon EOS Rebel T1i camera with a MP-E 65mm f/2.8 1-5x Macro Lens (Canon USA, New York, NY) mounted on a StackShot automated macro rail (Cognisys Inc., Traverse City, MI), and placed it into an individual cup. Individual chambers in each gall were visible as light green subcircles, and, especially in later collections, dark holes indicated that a galler, parasitoid, or inquiline had emerged (Figure 2A). Twice daily, we checked galls for a) emergent animals in cups, b) appearance of new emergence holes, c) signs of active chewing / movement at the gall surface, d) apparent head plugs, and e) previously identified head plugs that had been chewed through or otherwise destroyed. Head plugs were usually obvious, and defined as when gallers had chewed smaller-than-normal holes and had then stopped moving. We softly poked putative head plugs with the blunt end of a 0.20 mm minuten pin (BioQuip, Rancho Dominquez, CA) and only recorded an observation as a head plug when the insect made no movement in response. All observations were recorded daily on printed black and white photographs of each gall (Figure 2B). Late in the study, we dissected a subset of remaining head plugs to capture details of *Euderus* interactions with its host while inside the gall. In two chambers, we found gall wasps along with an apparently dead larva (Supplemental Figure 3). We extracted DNA from these larvae and sequenced mtCOI as above.

**Figure 2.**
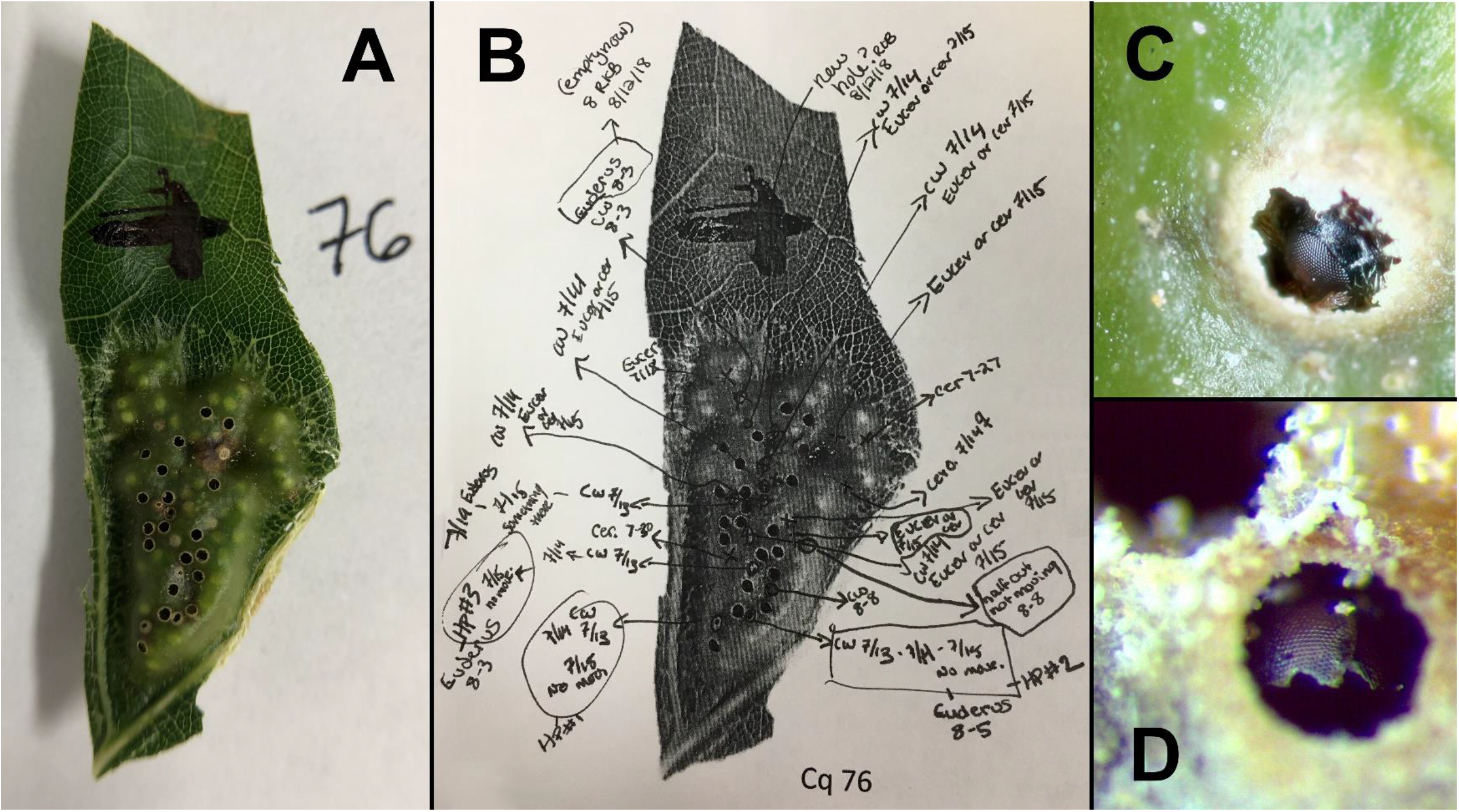
Details of methods for the *C. quercusmodesta / Euderus* emergence study. A) Photograph of a *C. quercusmodesta* gall one hour after collection, showing both intact chambers (light green circles) and galls from which an insect had emerged prior to the gall’s collection (dark circles). Very light tan-colored circles with a small pinprick of black in the center are animals that were either actively chewing out of their gall or had stopped chewing and were already “head plugs.” B) Example of notations made across the course of the study. Any changes to the gall were noted daily, such that all emergent animals found in the cup on any given day could be associated with the individual chamber from which they emerged. This particular gall had four *Euderus* emerge from gall chambers for which chewing (“CW”) and/or a “head plug” (“HP”) had previously been observed. Other notation refers to dates of observations, initials of observer (e.g., “RKB”), or the genus of the emergent animal (e.g., “Eucer” for the inquiline *Euceroptres*); C) close-up of a “head plug” where a *C. quercusmodesta* galler has chewed a partial hole and then stopped moving; D) close-up of *C. quercusmodesta* head after emergence of a *Euderus* parasitoid.

For the five other *non-Bassettia* gall wasp species from which *Euderus* emerged in the lab, we inspected post-emergence galls for evidence of head plugs, chewed heads, or other signs of hypermanipulation previously described in the *B. pallida* / *E. set* interaction [17].

## 3. Results

### (a) Collections and discovery of new Euderus / galler associations

Between 2015 and 2018 we collected more than 23,000 galls representing approximately 100 oak gall wasp species (Supplementary Table 1) and subsequently reared >15,000 individual parasitoids, inquilines, and hyperparasitoids (A.K.G.W. & A.A.F., unpublished data). Among these collections, we reared *Euderus* wasps from six different gall wasp host species (Table 1): *Andricus quercuspetiocola, Callirhytis flavipes, Callirhtyis quercusmodesta, Callirhytis quercussctiula, Callirhytis tumifica* and *Neuroteras noxiosus*. All *Euderus* keyed morphologically to *Euderus set*.

**Table 1.** Description of oak gall wasps, tree habitats, and geographic locations from which *Euderus set* has been reared and identified.

| Cynipid oak gall-wasp | Oak host(s) species and section | Locations | Description of gall | Timing of galls collections and *Euderus* emergences | Record |
| --- | --- | --- | --- | --- | --- |
| *Andricus quercuspetiolicola* | *Q. alba*;<br>*Q. bicolor*<br>(section: *Quercus*) | IA | Multichambered integral leaf gall, swelling of leaf petiole and sometimes basal midrib projecting on both sides of leaf | Galls collected May - August, most *E. set* emergences June - July | This study |
| *Bassettia pallida* | *Q. geminata*;<br>*Q. virginiana*<br>(section: *Quercus*) | FL, GA, LA, MS, TX | Integral stem gall (a “crypt”) | Galls collected July – October, most *E. set* emergences Feb – March. | [16, 17] |
| *Callirhytis flavipes* | *Q. macrocarpae*<br>(section: *Quercus*) | IA | Integral leaf gall, elongated swelling of the midrib | Galls collected May -July, most *Euderus* emergences June-July | This study |
| *Callirhytis quercusmodesta* | *Q. palustris*<br>*Q. rubra*<br>(Section: *Lobatae*) | IA, MO | Integral leaf gall, parenchyma of leaf swelling, projecting on both sides, multiple chambers | Galls collected May - August, most *E. set* emergences July-August | This study |
| *Callirhytis quercusscitula* | *Q. imbricaria*<br>(Section: *Lobatae*) | IA, MO | Swelling of stem at the base of where new leaves attach | Galls collected June, most *E. set* emergences June – July | This study |
| *Callirhytis tumifica* | *Q. coccinea*<br>(Section: *Lobatae*) | PA | Integral leaf gall, swelling of midrib on the bottom third of leaf | Galls collected May- June, most *E. set* emergences July | This study |
| *Neuroterus noxiosus* | *Q. bicolor*<br>(section: *Quercus*) | IA | Integral leaf gall | Galls collected June, most *E. set* emergences June-July | This study |

### (b) MtCOI sequencing

COI sequences also suggest that all *Euderus* wasps in this study were *E. set*. The 16 adult and two larval *Euderus* sequenced in this study had mtCOI sequences that were 95-100% identical to one another, differing by a maximum of 29 bp across the full 701bp sequence (Supplemental Figure 1). The four shorter sequences from 2016 differed by 0-21 bp (96 - 100% similarity) across their 461bp overlap with the other sequences. Twelve wasps had mtCOI that differed by just 6-15 bp (98% - 99% similarity) from the two sequences from *E. set* associated with *B. pallida*. Four other *Euderus* had mtCOI sequences that were somewhat more divergent (20-29 bp, 95 - 97% similarity) from the other wasps, but percentage sequence similarity should not alone be used to define species [25], and a lack of perceptible morphological differences and representation of the same gall wasp hosts in both haplotype groups both suggest that these are two somewhat divergent haplotype variants in a single species.

### (c) Observations of Euderus behavior and emergence

Across 128 *Callirhytis quercusmodesta* galls, we reared 291 adult *C. quercusmodesta* gallers, 44 *Euderus* wasps, and 649 other parasitoids and inquilines (Table 2; Supplemental Figure 2). During the course of the study we observed 63 *C. quercusmodesta* “head plugs.” Thirty-nine *Euderus* were conclusively linked to a specific head plug that had been noted during a previous observation period. The 24 head plugs from which no *Euderus* emerged did not produce any other adult parasitoids, suggesting that *Euderus* died inside of the chamber as in the two we found during dissections (Supplemental Figure 3). In five cases, a *Euderus* emerged from a gall where we had not previously observed a head plug, indicating either a failure of detection or a genuine lack of a plug. Evidence of head-plugs, both intact and eviscerated, was also found in all other gall collections from which *Euderus* wasps emerged (Figure 3).

**Table 2.** Insects reared from 128 *Callirhytis quercusmodesta* leaf gall clusters from a single Pin oak (*Quercuspalustris*) in Iowa City, IA.

| Family | Genus | No. reared |
| --- | --- | --- |
| Eupelmidae | *Brasema* | 12 |
| Eulophidae | *Euderus* (*E. set*) | 44 |
|  | *Galeopsomyia* | 30 |
| Eurytomidae | *Eurytoma* | 75 |
|  | *Sycophila* (3 species) | 82 |
| Ormyridae | *Ormyrus* | 5 |
| Platygastridae | unknown | 8 |
| Cynipidae | *Ceroptres* | 426 |
|  | *Callirhytis* (galler) | 291 |
| Cecidomyiidae<br>(Diptera) | unknown | 11 |

**Figure 3.**
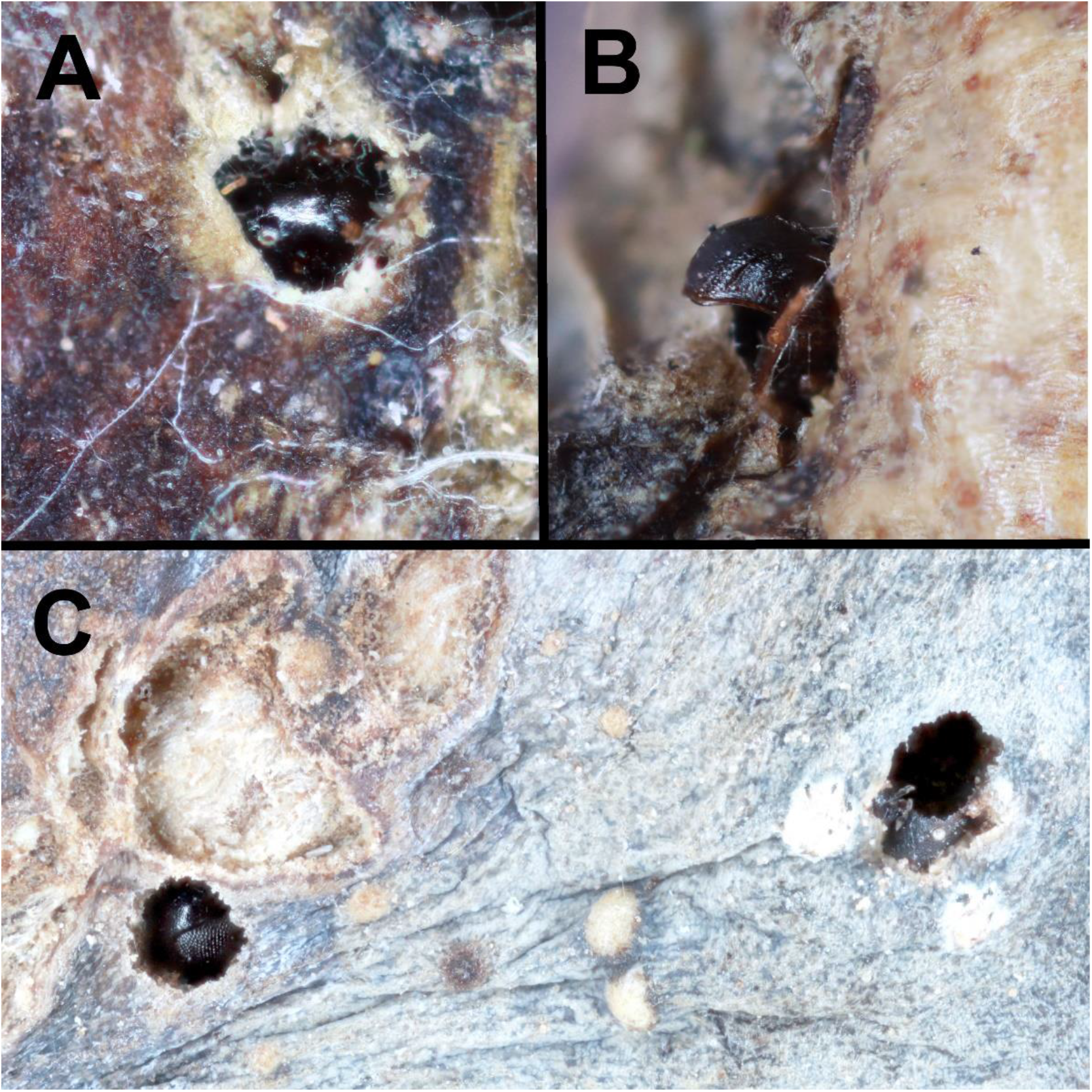
Evidence of *Euderus set* activity in galls formed by three other cynipid gall wasps in oaks. A) Head plug of *Callirhytis quercusscitula* in a gall; B) *Callirhytis tumifica* mesothorax (head missing) extruding from an exit hole. This phenotype was also seen in *B. pallida* galls studied in Egan et al. [16]; C) *Andricus quercuspetiolicola* head plug with eye visible (left) and partially chewed *A. quercuspetiolicola* head post-*E. set* emergence (right).

## 4. Discussion

Questions regarding the evolution of ecological specialization have vexed biologists for millennia [1]. Why do many taxa evolve to use only a subset of resources available to them in the environment? This general pattern of specialization is thought to be influenced by trade-offs, where specialization on one resource is maladaptive to others. For host-parasite systems that involve behavioral manipulation, pattern and process have not been explored in great detail. In part, this is due to a dearth of study systems available to address these important questions.

### (a) Euderus *in North America*

Morphological and genetic data together imply that all *Euderus* wasps reared in this study are *E. set*, the species described previously from *Bassettia pallida* [16], bringing its total number of confirmed hosts to seven. *Euderus set* remains the only endemic cynipid-associated *Euderus* known from the eastern half of North America. In Arizona and California, *Euderus crawfordii* has been reared from the native oak gall wasp *Dryocosmus coxii* on Emory Oak *(Quercus emoryi)* and Silverleaf Oak (*Quercus hypoleucoides*) [18,26], as well as from the introduced gall wasp *Plagiotrochus suberi* on non-native Cork Oak (*Quercus suber*) [27,28]. While we have no sequence data available for this southwestern species, the morphological resemblance [16] and similarity of habit between *E. crawfordii* and *E. set* suggest they may be sister to one another. Another possible cynipid parasitoid, *Euderus albitarsis*, was introduced to the Eastern U.S. for control of the western larch case bearer (*Coleophora laricella* [Lepidoptera: Coleophoridae]) [26], but the single reference to an association with an unknown cynipid [29] has no independent corroboration, so this record may be aberrant.

The current study does not allow for a complete accounting of *E. set’*s host range across seasons and geographic distribution. Previous work described an overwintering habitat for *E. set* in a southern twig-galling wasp but did not identify a spring or summer host [16]. Conversely, here we do not identify an overwintering host for *E. set* in any of the more northerly locations from which it was reared, though our collections did include several overwintering galls (e.g., *Andricus dimorphus, Andricus quercusstrobilanus, Dryocosmus imbricarae*).

### (b) Host specificity in E. set

We find that *Euderus set* manipulates at least seven host species spanning four genera (Table 1). While some parasites are more effective at manipulating more closely related hosts [e.g., 30], the ability to effectively manipulate the behavior of distantly related hosts is not unknown. For example, the parasitoid wasp *Zatypota kauros* can alter the web-building behavior of spider hosts from different families [31]. It is also worth noting that the hosts of *E. set* may not be as distantly related as their names suggest. The taxonomy of gall wasps is fluid, with taxa switching among genera with some regularity [26,32,33]. Indeed, the gall wasp genera *Bassettia* and *Callirhytis* are morphologically similar [34], to the extent that some species have been indeterminately placed in one genus versus the other [e.g., 35]. On the other hand, *Andricus quercuspetiocola* and *Neuroterus noxiosus* are seemingly more distantly related, at least based on morphological characters [32]. At best, we might say that some – but not all – of the known hosts of *E. set* may be more closely related than their specific names imply, but without a comprehensive phylogeny of North American gall wasps, these conclusions remain tentative.

Many potential gall wasp hosts were also collected from the same tree host species at the same time as *E. set*, and yet were not apparently manipulated or even infected by *E. set*. Our collections (Supplementary Table 1) represent approximately 100 of the ~700 described species of Nearctic oak gall wasps [36]. If relatedness of hosts determined host range for *E. set*, we would expect that other *Callirhytis* in our collections might have yielded *Euderus*, especially those that we collected in large numbers (e.g., *C. punctata, C. futilis, C. quercusgemmaria)*. So while *E. set* is oligophagous and widespread (Iowa to Pennsylvania; south to Texas and Florida), it is also not a broad generalist on all oak gall wasps.

If *E. set* is able to manipulate gall wasps from disparate genera residing on multiple different host plant species, then why do we not find *E. set* manipulating *all* gall wasps from the host plants on which we find this parasitoid? First, in some cases, it is reasonable to assume that we did not rear *E. set* from some hosts because it has a patchy distribution, and/or because our collection numbers were variable from one gall species to another. Negative rearing results should always be interpreted with caution. On the other hand, it is also possible that something other than host phylogeny constrains *E. set’s* ability to infect and manipulate particular hosts. The presence of behavioral alterations in multiple cockroach species infected by the acanthocephalan *Moniliformis moniliformis* was not predicted by host phylogeny [37]. Additionally, the brain-infecting trematode *Euhaplorchis sp. A* is able to infect and manipulate particular fish from two families (Fundulidae and Poecilidae), but is not able to infect and manipulate all fish in these families (i.e., it could not infect *Lucania parva*, a fish in the Fundulidae family) [38]. In both of these studies, controlled infections confirmed that parasites given access to a variety of hosts were unable to infect and/or manipulate a subset of these hosts, suggesting that something other than phylogeny is important in determining host specificity.

In this study we did not employ controlled infections, so we cannot be sure *E. set* had an opportunity to encounter each host type. However, we never observed a possible gall wasp host species from which *E. set* emerged in the absence of manipulation (e.g., presence of head plugs), nor did we observe *E. set* dead inside of the galls of any other species. This argues against the idea that there are hosts that *E. set* is able to successfully parasitize but not manipulate. Thus, the strongest emergent hypothesis for what limits host range for *E. set* is not the identity of the gall wasp host, but the physical characters of the gall itself. Authors have described *Euderus* as larval or pupal parasitoids [18,39], and they lack the long, exerted ovipositors typical of some other genera of late-attacking galler parasitoids (e.g., genus *Torymus* [40]), such that they may be limited to attacking hosts that are not buried deep within a gall nor protected by complex defense structures. The galls of all seven known hosts are integral (i.e., enclosed within the epidermis; not detachable without causing significant damage to the plant tissue), such that when a developing gall wasp approaches the pupal stage and fills its gall chamber, it will be near the surface and physically accessible to *E. set*. All known hosts also lack the baroque structural defenses found on many other galls, which can include spines, fuzz, or larval cells suspended deep inside otherwise empty chambers [41]. These defenses grow more substantial as the gall grows, which may render many galls inaccessible to *Euderus* and other parasitoids of gall wasp pupae.

The phenology of the *C. quercusmodesta* system also bears out the narrative that *E. set* specializes on a subset of relatively unprotected gall wasp pupae. Though we collected *C. quercusmodesta* galls weekly across six weeks, *E. set* emerged only from galls collected in weeks four through six (7/6, 7/13/ & 7/20), corresponding with the timing of *C. quercusmodesta* pupation and adult emergence (Supplementary Figure 4). This supports the idea that *E. set* attacks galls close to when the gall wasp is mature and ready to emerge from the gall. Further, this temporal pattern of attack is not due to mature *E. set* being scarce in the environment until early July: emergence data from other hosts (Supplemental Figure 5) show that adult *E. set* are present in nature well before *C. quercusmodesta* galls mature. Other authors have also suggested that host ecology may play an important role in determining which hosts are manipulated by parasites [e.g., 38], but thus far no studies have addressed this question explicitly.

Parasites that manipulate multiple host species do not always induce the same manipulated phenotypes in all host species (e.g., [35]). However, the observation that *E. set* induces the same behavior in multiple hosts suggests that it is exploiting a mechanism conserved across all of the manipulated hosts. At this time we can only speculate on what this shared mechanism would be. Fredensborg’s study [9] suggested that parasites with distantly related hosts are more likely to use debilitation to manipulate these hosts.

This suggests that the “head-plugging” behavior may arise by debilitating the host at an appropriate time. But how does *E. set* “know” when to debilitate the host? Since we know from the *C. quercusmodesta* collections that attack occurs before the chewing of the exit hole (some holes were not chewed until galls were brought into the lab), some signal must induce *E. set* to interrupt the normal behavior of the parasitized gall wasp. Could this signal be the production of cellulase, which some insects use to digest plant material? Alternatively, perhaps the trigger is a plant-related signal, produced exclusively by the epidermal tissue and encountered only as the gall wasp begins its exit, or a response by *E. set* to an increase in light as its host chews its exit hole. The specific mechanisms underlying host control and the rules underlying host range promise rich foundations for future research in this system.

### (c) Evolution of crypt-keeping

When and how did this behavior evolve? Though very little is known about the biology and behavior of most *Euderus* wasps, a study of *Euderus lividus*, parasitoid of *Agromyza obtusa* [Diptera: Tephritidae] suggests what may be a precursor to this system, though without apparent behavioral manipulation. Larval *A. obtusa* chew an exit hole in the side of the seed pods in which they are feeding. Ahmad [42] found several new exit holes containing incapacitated *A. obtusa* larvae alongside *E. lividus* eggs, suggesting that attack had occurred soon after the exit hole had been chewed. The exoparasitic *E. lividus* larvae then fed on the *A. obtusa*, after which adult parasitoids developed and exited the holes. Ahmed [42] further suggested that without the exit hole, *E. lividus* would have been trapped in the now extremely hard seed pod. In the case of *E. set*, we know that attack can occur before the chewing of the exit hole, because in our *C. quercusmodesta* study some holes that became head plugs were chewed by adult gall wasps after galls were brought in from the field. If other *Euderus* species are also found to manipulate host behavior, a phylogeny of *Euderus* that maps the presence, absence, and species-specific nature of behavioral manipulation may aid in understanding the evolution of this trait.

### (d) Synthesis and conclusions

There is a near universal pattern of parasites to be highly host specialized. For parasites that manipulate the behavior of their hosts, the symbiotic intimacy implied by behavioral control might be expected to further restrict host range – though the literature to date is equivocal on this point [9, 37, 38]. We have discovered that the behavior-manipulating parasitoid wasp *E. set* is specialized on a subset of available gall wasp hosts, but that these hosts are taxonomically diverse. Remarkably, each of *E. set’s* hosts has converged upon similarities in their extended phenotypes suggesting that behavioral manipulation may be less important to its host range than other dimensions of the host-parasitoid interaction, such as host’s physical defenses.

## Data accessibility

All data are available as electronic supplementary material.

## Authors’ contributions

A.K.G.W. and A.A.F. conceived of the study. A.K.G.W. and O.S.K. conducted the *C. quercusmodesta* study. All authors contributed to gall collections, insect rearing, and writing of this paper.

## Acknowledgements

Robin Bagley, Will Carr, Sara Devine, Rachel Erikson, Leo Gastel, Alaine Hippee, Emily Manders, Danny McGarry, Moe Shakally, Joseph Verry, and Caleb Wilson assisted with gall collections and rearing and preservation of insects. Thanks to Robbins Park Environmental Education Center in Ambler, PA for permission to collect *C. tumifica* galls from a scarlet oak on their property. Figure 1 was illustrated by the incomparable Mona Luo (http://www.monaluo.com/).

