## Supplemental Table 2 and figures 1-5 for "A keeper of many crypts: the parasitoid *Euderus set* manipulates the behavior of a taxonomically diverse array of oak gall wasp species"

Supplementary materials

Supplemental Table 2. Species, host, tree, and collection site information for twenty *Euderus set* wasps analyzed in this study. The two *E. set* reared from *Bassettia pallida* were previously reported in Egan (2017).

| Genbank accession # | Species | Gall-wasp host | Tree host | Location | Primers | bp length | Lab specific ID# |
|---|---|---|---|---|---|---|---|
| MK294992 | *Euderus set* | *Neuroterus noxiosus* | *Quercus bicolor* | SA:Iowa City,IA | COI_pf2/COI_2437d | 664 | 341-053-38 |
| MK294993 | *Euderus set* | *Neuroterus noxiosus* | *Quercus bicolor* | USA:Iowa City,IA | COI_pf2/COI_2437d | 635 | 369-053-6 |
| MK294994 | *Euderus set* | *Callirhytis flavipes* | *Quercus macrocarpae* | USA:Tiffin,IA | COI_pf2/COI_2437d | 701 | 919-013-5 |
| MK294995 | *Euderus set* | *Andricus quercuspeticola* | *Quercus alba* | USA:Tiffin,IA | COI_pf2/COI_2437d | 675 | 920-010-12a |
| MK294996 | *Euderus set* | *Andricus quercuspeticola* | *Quercus alba* | USA:Tiffin,IA | COI_pf2/COI_2437d | 636 | 920-010-16a |
| MK294997 | *Euderus set* | *Andricus quercuspeticola* | *Quercus bicolor* | USA:Iowa City, IA | COI_pf2/COI_2437d | 701 | 949-042-9 |
| MK294998 | *Euderus set* | *Callirhytis quercusscitula* | *Quercus imbricaria* | USA:Creve Coeur, MO | COI_pf2/COI_2437d | 685 | 1409-049-4 |
| MK294999 | *Euderus set* | *Callirhytis tumifica* | *Quercus coccinea* | USA:Ambler,PA | COI_pf2/COI_2437d | 664 | 1416-043-4 |
| MK295000 | *Euderus set* | *Callirhytis quercusmodesta* | *Querucs palustris* | USA:Iowa City,IA | LepF/LepR | 652 | 20-1-5c |
| MK295001 | *Euderus set* | *Andricus quercuspeticola* | *Quercus macrocarpae* | USA:Tiffin,IA | COI_pf2/COI_2437d | 701 | 918-042-19a |
| MK295002 | *Euderus set* | *Callirhytis flavipes* | *Quercus macrocarpae* | USA:Tiffin,IA | COI_pf2/COI_2437d | 701 | 919-013-4a |
| MK295003 | *Euderus set* | *Callirhytis quercusscitula* | *Quercus imbricaria* | USA:Creve Coer,MO | COI_pf2/COI_2437d | 701 | 1409-049-15 |
| MK295004 | *Euderus set* | *Callirhytis tumifica* | *Quercus coccinea* | USA:Ambler,PA | COI_pf2/COI_2437d | 701 | 1416-043-1 |
| MK295005 | *Euderus set* | *Callirhytis quercusmodesta* | *Quercus palustris* | USA:Iowa City,IA | COI_pf2/COI_2437d | 655 | cq94 |
| MK295006 | *Euderus set* | *Callirhytis quercusmodesta* | *Quercus palustris* | USA:Iowa City,IA | COI_pf2/COI_2437d | 701 | cq115 |
| MK295007 | *Euderus set* | *Callirhytis quercusmodesta* | *Quercus palustris* | USA:Iowa City,IA | LepF/LepR | 652 | 20-1-5B |
| MK295008 | *Euderus set* | *Callirhytis quercusmodesta* | *Quercus palustris* | USA:Iowa City,IA | LepF/LepR | 420 | 17-1-11A |
| MK295009 | *Euderus set* | *Callirhytis quercusmodesta* | *Quercus palustris* | USA:Iowa City,IA | LepF/LepR | 652 | 17-1-9C |
| MK295010 | *Euderus set* | *Bassettia pallida* | *Quercus geminata* | USA: Inlet Beach, FL | COI_pf2/COI_2437d | 703 | Es_1 |
| MK295011 | *Euderus set* | *Bassettia pallida* | *Quercus geminata* | USA: Inlet Beach, FL | COI_pf2/COI_2437d | 744 | Es_2 |

Supplemental Figure 1. Bayesian network of mtCOI sequences for *Euderus* samples. Branches are marked with posterior probabilities. The two *Euderus set* samples from Egan et al. [16] are in black bold font. Other *Euderus set* sequences are color-coded based on host association: Blue = *Callirhytis quercusscitula*; Red = *Callirhytis tumifica*; Purple = *Callirhytis quercusmodesta*; Green = *Callirhytis flavipes*, Yellow = *Andricus quercuspeticola*; Dark blue = *Neuroterus noxiosus*. Tree host associations are indicated by the following abbreviations: Qa = *Quercus alba*, Qb = *Quercus bicolor*, Qc = *Quercus coccinea*, Qg = *Quercus gemmaria*, Qi = *Quercus imbricarae*, Qm = *Quercus macrocarpae*, Qp = *Quercus palustris*. Location of collections are indicated by state abbreviation. For additional details, see Supplemental Table 2.

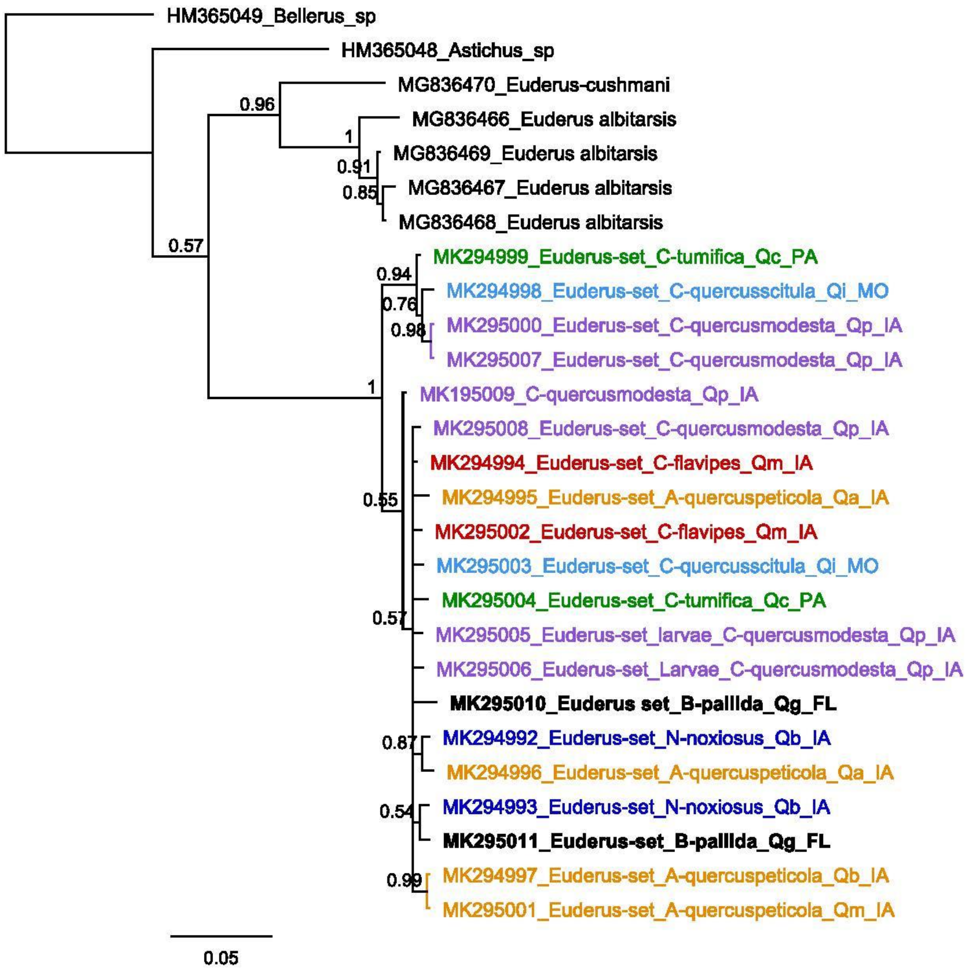

Supplemental Figure 2. Eclosion timing of *C. quercusmodesta* and seven of its hymenopteran gall associates, including *E. set*.

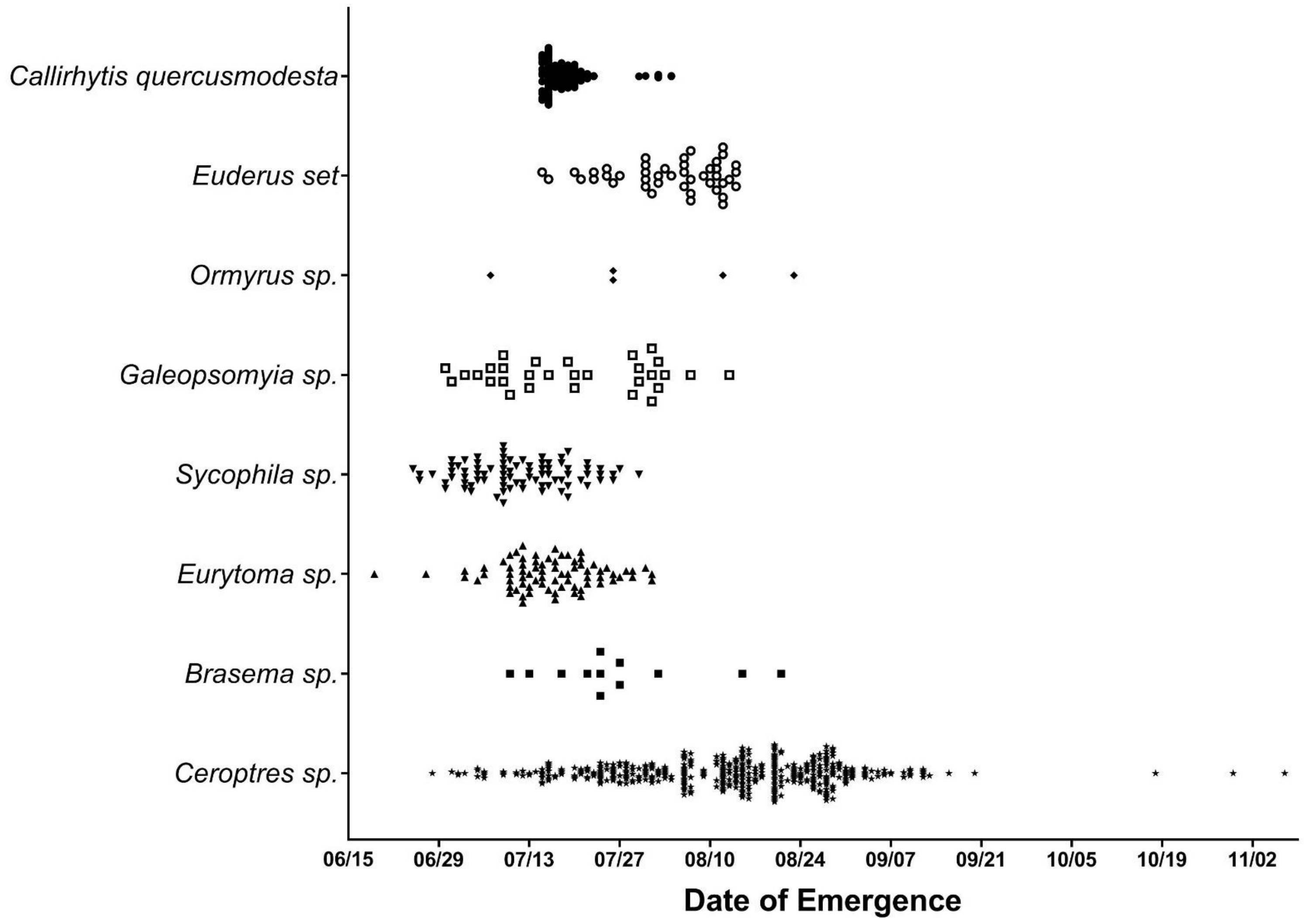

Supplemental Figure 3. Cross section (side view) of a head plug from a *Callirhytis quercusmodesta* gall. The adult gall wasp is towards the top of the image with its head plugging a partially chewed exit hole. Beneath the gall wasp (arrow) a larval *Euderus* is visible. DNA extraction and sequencing of mtCOI showed this larva to be *Euderus set*.

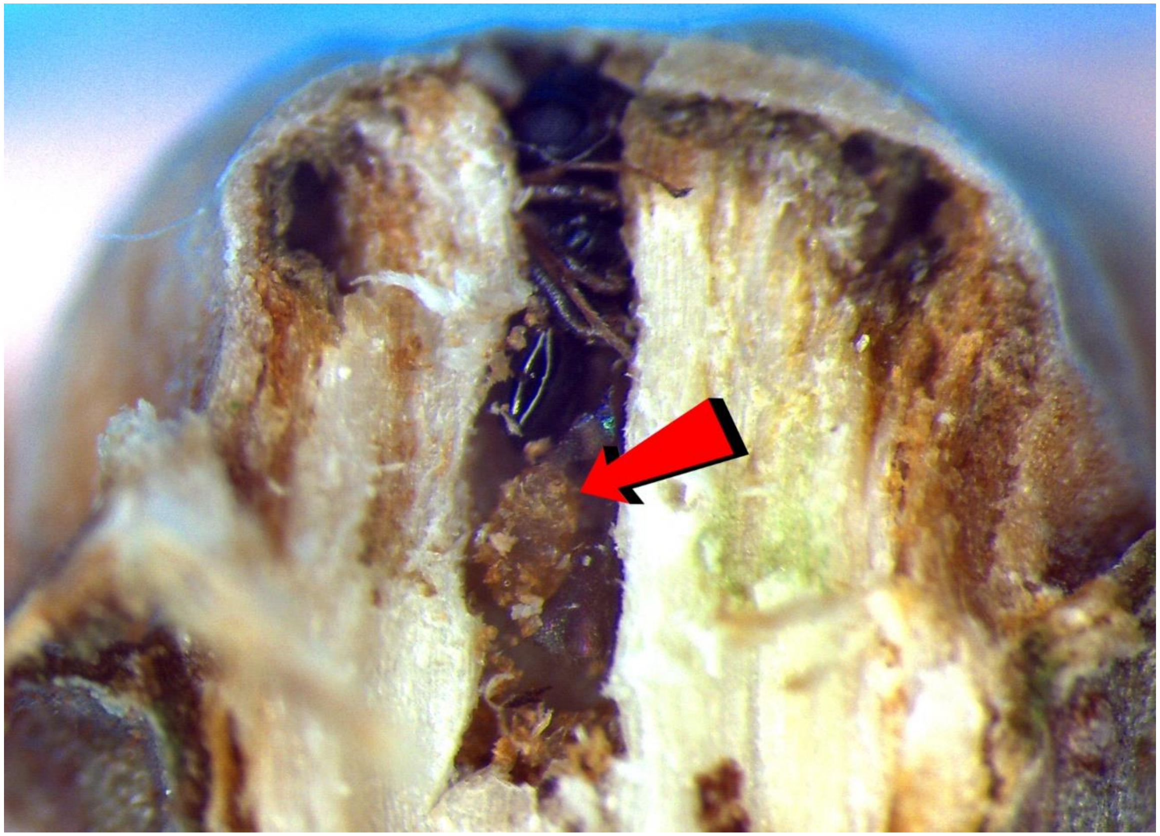

Supplemental Figure 4. Emergence of *E. set* (open circles) from six collections of *C. quercusmodesta* galls in 2018. Dotted vertical lines reference the dates of collections. Unparasitized *C. quercusmodesta* (closed circles) emerged from all six collections, but all emerged after the date of the fifth collection (7/13).

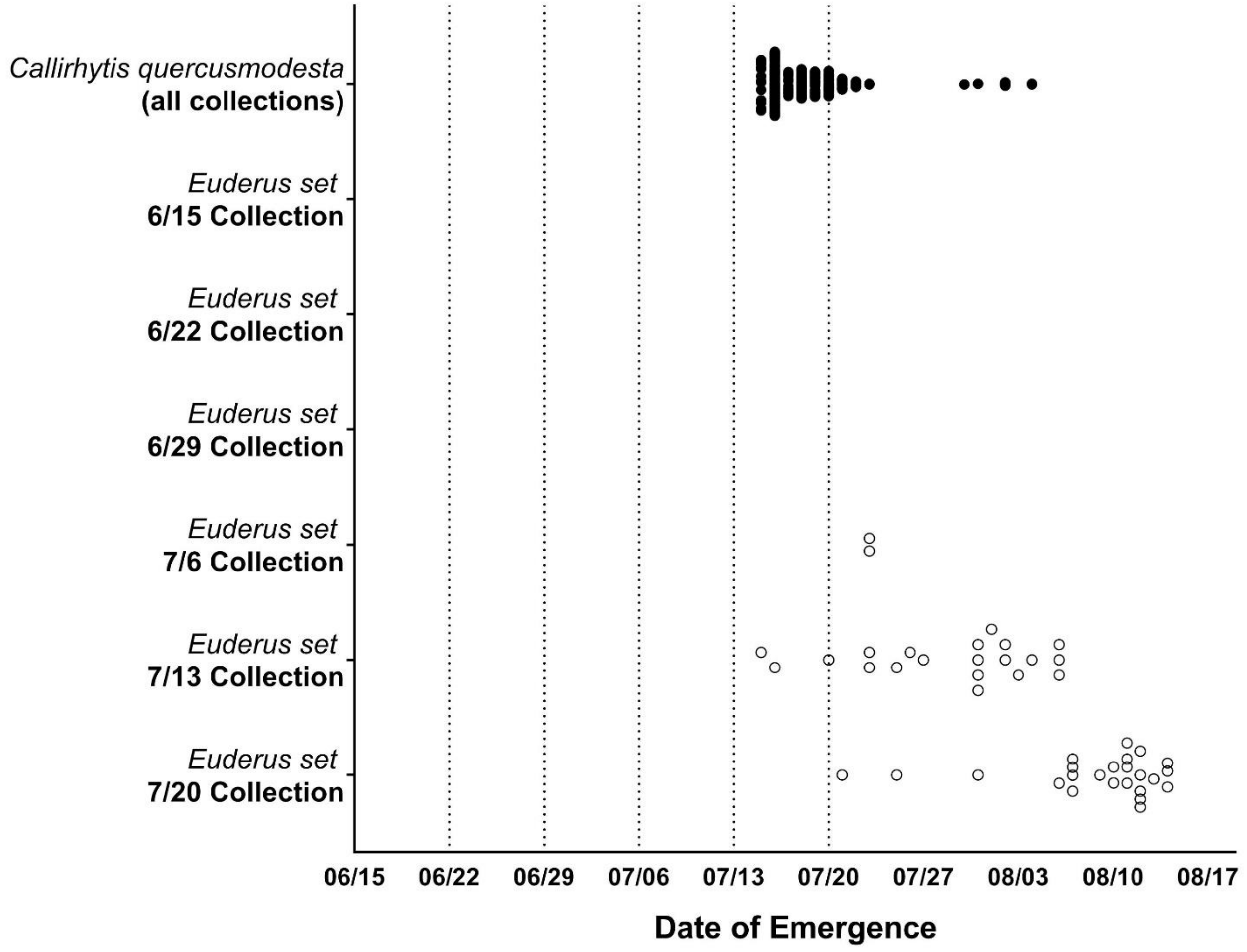

Supplemental Figure 5. Emergence phenology of gallers (closed circles) and their associated *E. set* (open circles) from galls on six oak tree species. Tree habitat abbreviations: Qb = *Quercus bicolor*; Qi = *Quercus imbricaria*; Qc = *Quercus coccinea*; Qa = *Quercus alba*; Qm = *Quercus macrocarpae*; Qp = *Quercus palustris*. Galler host abbreviations: Nn = *Neuroterus noxiosus*; Cqs = *Callirhytis quercusscitula*; Ct = *Callirhytis tumifica*; Aq = *Andricus quercuspetiolicola*; Cf = *Callirhytis flavipes*; Cqm = *Callirhytis quercusmodesta*.

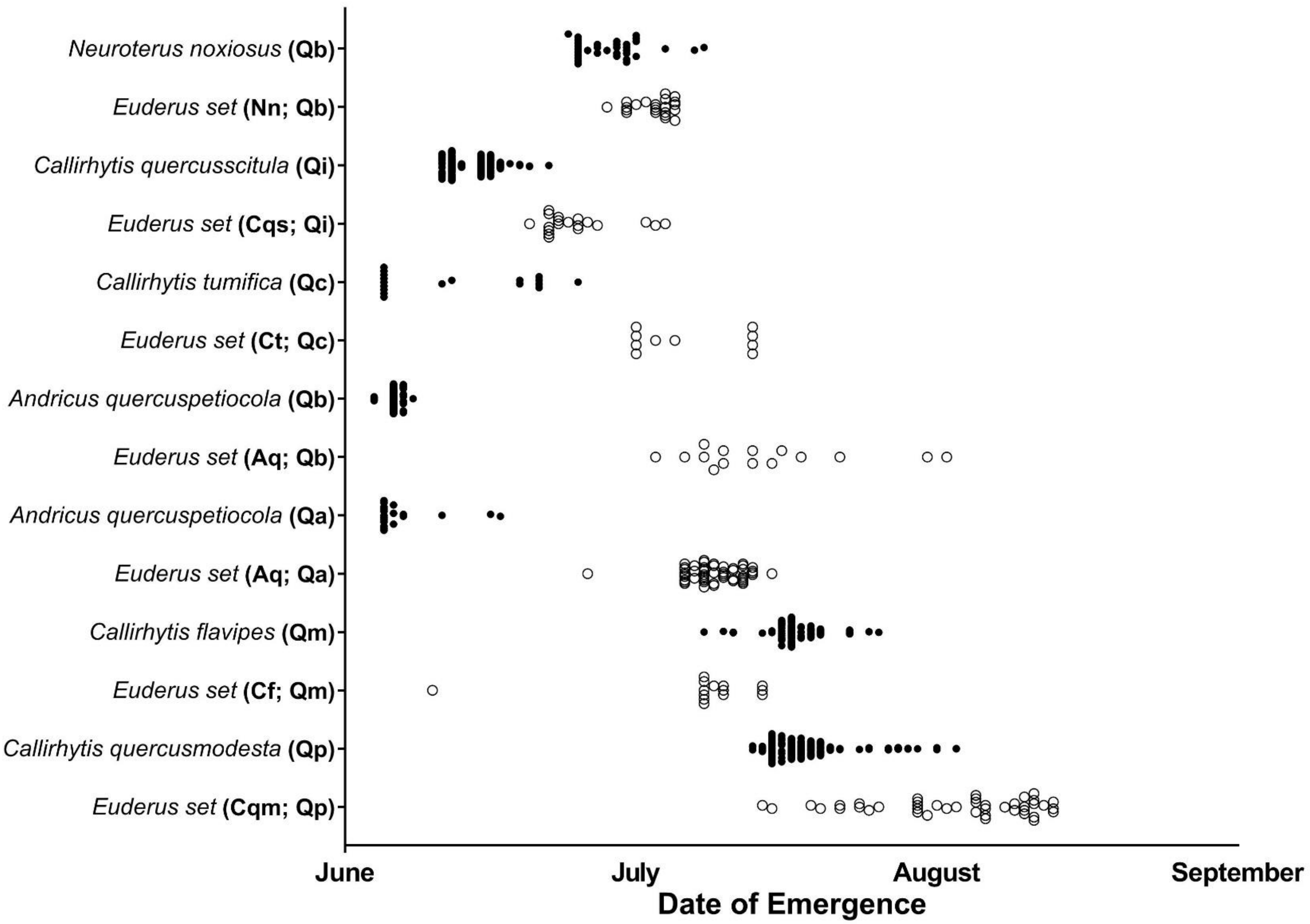
